# Leg force interference in poly-pedal locomotion

**DOI:** 10.1101/269100

**Authors:** Tom Weihmann

## Abstract

The examination of gaits and gait-changes have been the focus of movement physiology and legged robot engineering since the first emergence of the fields. While most examinations focussed on bipedal and quadrupedal designs many robotic implementations rely on the higher static stability of three or more pairs of legs. Nevertheless, examinations of gait-changes in the biological models, i.e. poly-pedal arthropods such as insects or arachnids, are rare. Except for the well-known change from slow feedback controlled walking to a fast, feedforward controlled running gait, no changes are known or are deemed to be of low significance.

However, recent studies in fast moving spiders, mites and cockroaches have revealed an additional gait change also for the transition from intermediate to high running speeds. This change is similar to gait transitions as found in quadrupedal vertebrates.

Accordingly, the present approach aims to extend available theory to poly-pedal designs and examines how the number of active walking legs affects body dynamics when combined with changing duty factors and phase relations. The model shows that higher numbers of active leg pairs can prevent effective use of bouncing gaits such as trot and their associated advantages because significantly higher degrees of leg synchronisation are required. It also shows that small changes in the leg coordination pattern have a much higher impact onto the COM dynamics than in locomotor systems with fewer legs. In this way, the model reveals coordinative constraints for specific gaits facilitating the assessment of animal locomotion and economization of robotic locomotion.

**Significance Statement:** The present model approach enables to assess the impact of different numbers of walking legs onto movement dynamics and gait choice in terrestrial legged locomotion. The model’s results are indicatory for research in legged locomotion regardless whether biological examples or legged walking machines are considered. The approach is suitable for all numbers of pairs of walking legs larger than two and is focussed on symmetrical gaits as found in straight and continuous locomotion. The model fills a gaping gap as the impact of phase shifts among the legs in the coordinated sets of legs typical for poly-pedal animals and robots on overall body dynamics are not considered sufficiently in existing dynamic model approaches.

## Background

Usually intermediate and fast legged locomotion in terrestrial arthropods such as insects and arachnids is considered as closely connected to the synchronous activity of diagonally adjacent legs, for instance tripodal sets of legs in insects (Hughes, 1952; Wilson, 1966). By contrast, recent research revealed deviating symmetrical (Hildebrand, 1965) leg coordination patterns in a couple of fast running species such as cockroaches, mites and spiders (Weihmann, 2013; Weihmann et al., 2017; Weihmann et al., 2015). Given appropriate leg properties, alternating sets of legs facilitate the employment of spring mass dynamics well known from running bipeds and trotting terrestrial vertebrates (Alexander, 1989; Blickhan and Full, 1993). Locomotor apparatuses with legs acting similarly to mechanical springs that support the body during stance are found to be energy efficient structures which provide high degrees of dynamic stability (Blickhan and Full, 1993; Full and Koditschek, 1999).

Locomotor apparatuses rely on the mechanical and physiological properties of their components, i.e. the legs which constitute it. In arthropods moving straight ahead, frontal leg pairs only decelerate and rear legs only accelerate (Full et al., 1991; Reinhardt et al., 2009; Weihmann and Blickhan, 2006). Accordingly, most arthropod legs experience either flexion or extension during a contact phase which effectively prevents energy storage and recovery within the legs. The only structures that can store energy are the hips if they were first deflected in dorsal direction and flex ventrally in the later stance phase (Dudek and Full, 2006). This implies that only vertical forces can be recovered and vertical amplitudes of the COM directly affect the capacity of elastic energy storage. This relation is less significant in fast moving vertebrates with their flexible spines and erect legs. In vertebrates, all legs can contribute to deceleration and acceleration (Robin et al., 2008); each leg can store movement energy in the initial stance phase and recycle it in the later stance phase. Accordingly, the vertical amplitude of the COM is not necessarily equivalent to length changes of the leg springs in vertebrates and the amount of temporarily stored elastic energy (Farley et al., 1993); though the stiffness per leg appears to be similar among a wide range of species and body plans (Blickhan and Full, 1993).

However, with increasing running speed and declining contact durations, spring mass dynamics require increasing vertical amplitudes of the centre of mass (COM) (Blickhan, 1989; McMahon and Cheng, 1990) which might be hampered by the typically sprawled legs and the low COM positions in arthropods. If required vertical oscillations cannot be maintained either by limitations of the material properties of the load bearing skeletal components (Taylor and Dirks, 2012), the specific design or position of the legs (Günther and Weihmann, 2011; Seyfarth et al., 2001) or by limitations of the muscular system (Guschlbauer et al., 2007; Siebert et al., 2010) the elastic capabilities of tendons and other compliant structures can no longer be appropriately exploited.

If legs are considered as damped mechanical oscillators (Alexander, 1997; Dudek and Full, 2006; Rubenson et al., 2004) the ratio of potentially recovered and dissipated energy, i.e. the damping ratio, gains importance. While oscillation energy grows with the second power of the oscillation amplitude, energy dissipation relies on viscoelastic processes and increases with the second power of speed. The mean vertical speed of the COM in spring mass systems during stance depends on the amplitude and the available time frame, i.e. the contact period. Since contact periods decrease hyperbolically with running speed (Hoyt et al., 2000; Weihmann, 2013; Weihmann et al., 2017) the vertical speed of the COM increases exponentially even if amplitudes would remain constant. The dissipated energy grows with the second power of this speed, such that the slope of the dissipated energy becomes much steeper than that of the potentially recovered oscillation energy. Consequently, with increasing running speed and depending on the damping ratio, the locomotor system may transform from an underdamped to an overdamped oscillator, which eventually would prevent energy efficient oscillations. Small animals such as insects are particularly prone to high damping ratios (Ache and Matheson, 2013; Garcia et al., 2000; Hooper et al., 2009). Accordingly gait changes are to be expected if the animals pass through significant ranges of contact durations and running speeds, even though arthropods and most other animals maintain high running speeds only for short periods of time (Bramble and Lieberman, 2004; Full and Weinstein, 1992).

Mathematical models are widely used for the computation and analysis of legged locomotion in men, birds and to some extent even for quadrupedal and hexapedal animals and robotic implementations. However, reduced analytic and numerical model concepts, struggle with temporally overlapping activities of more than two legs or synchronised sets of legs (Geyer et al., 2006; Schmitt et al., 2002). In poly-pedal organisms with temporally shifted activities and varying degrees of overlap between successive stride phases, such lumped model approaches are not applicable. Kinematic models in turn are well suited for the description of the temporal activities of the single legs within locomotor apparatuses with arbitrary numbers of legs (Cruse et al., 2007; Hildebrand, 1965; Wendler, 1964; Wilson, 1966). However, these model approaches largely disregard locomotion dynamics and ground reaction forces exerted by the legs. Therefore, they don’t give any insight into the consequences of the legs’ activities on movement dynamics, or whether requirements for certain COM dynamics can be fulfilled by an animal.

Therefore, the present study, aims to combine the best of both worlds by introducing typical single leg ground forces to a kinematic model approach. Major outcomes of spring mass dynamics are the following: similar, approximately symmetrical vertical ground forces of all walking legs and summed up vertical forces that oscillate about body weight (*bw*) (Blickhan and Full, 1993; Full and Tu, 1990). Ground reaction forces induce changes in the momentum and kinetic energy of the animal’s body. Accordingly, oscillation amplitudes of an animal’s body are determined by those of overall ground reaction forces (Alexander and Jayes, 1978; Reinhardt et al., 2009). In particular, the vertical ground force and oscillation amplitudes of the COM are crucial for the efficacy of elastic structures in the walking legs with regard to the storage and recovery of movement energy (Dudek and Full, 2006; Günther and Weihmann, 2012).

The essential parameters in kinematic descriptions of gaits are the duty factor, i.e. stance period divided by stride period, and the phase shifts between ipsilateral adjacent legs θ (Hildebrand, 1989). Both measures directly affect the maximum total forces exerted by the legs and resulting COM dynamics.

The present approach provides insight into the effects of changing leg numbers, duty factors and θ on the fluctuation amplitudes and frequencies of total vertical ground reaction forces. It shows that maximum force amplitudes decline more quickly with increasing deviation from alternating leg coordination as the number of walking legs increases. Duty factors in turn dramatically affect the absolute force amplitudes but have no effect on the relative decline over the phase range. Phase shifts between adjacent ipsilateral walking legs deviating significantly from 0.5 (alternation) or 1 (concurrency) increase the minimum number of legs in contact with the ground, which presumably leads to increased proprioception and controllability of the locomotion.

## Results

Depending on the number of legs involved in the generation of ground reaction forces, even relatively small changes in θ may result in significant alteration of the vertical force amplitudes (Fig. 1). However, the absolute changes are small if duty factors are high, and become substantial if duty factors are lower than about 0.7 (Fig. 2). Independent from the number of legs, maximum amplitudes of vertical forces are about 1 *bw* with duty factors of 0.5. Since the vertical momentum generated by the walking legs has to compensate for bodyweight over a stride, peak-to-peak values (i.e. the difference between the lowest and maximum force values) are twice as high (Fig. 1E). Maximum force amplitudes are 1.65 *bw* with duty factors of 0.2 and decrease almost linearly with increasing duty factors (Fig. 2). At a duty factor of 0.8 maximum force amplitudes are 0.23 *bw*, which is only about 1/7th of the former value.

**Fig. 1.**
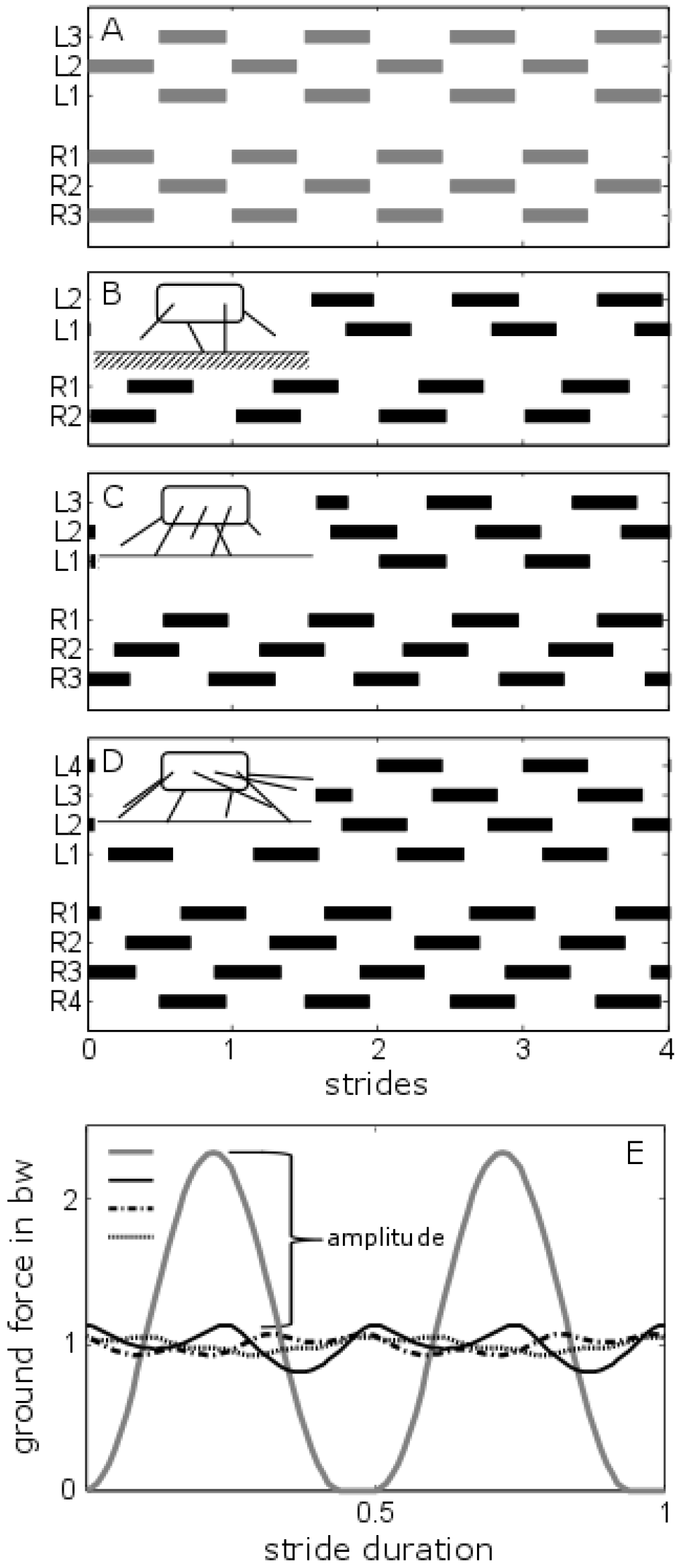
Gait patterns of four, six and eight legged locomotor systems (A-D) and corresponding overall ground forces over a stride (E) for single leg duty factors of 0.45. Bars represent contact phases; left (L) and right (R) legs are counted fore to aft. A) Gait pattern of a trotting hexapod; ipsilateral phases are 0.5.B-D Ipsilateral phase shifts (0.26, 0.34, 0.38) represent minima of vertical force amplitudes for the respective numbers of legs (4, 6, 8). These gait patterns correspond to the overall vertica ground forces shown in E) The grey solid line refers to A, the black solid line to B, the dash-dotted line to C and the dotted line to D.

**Fig. 2.**
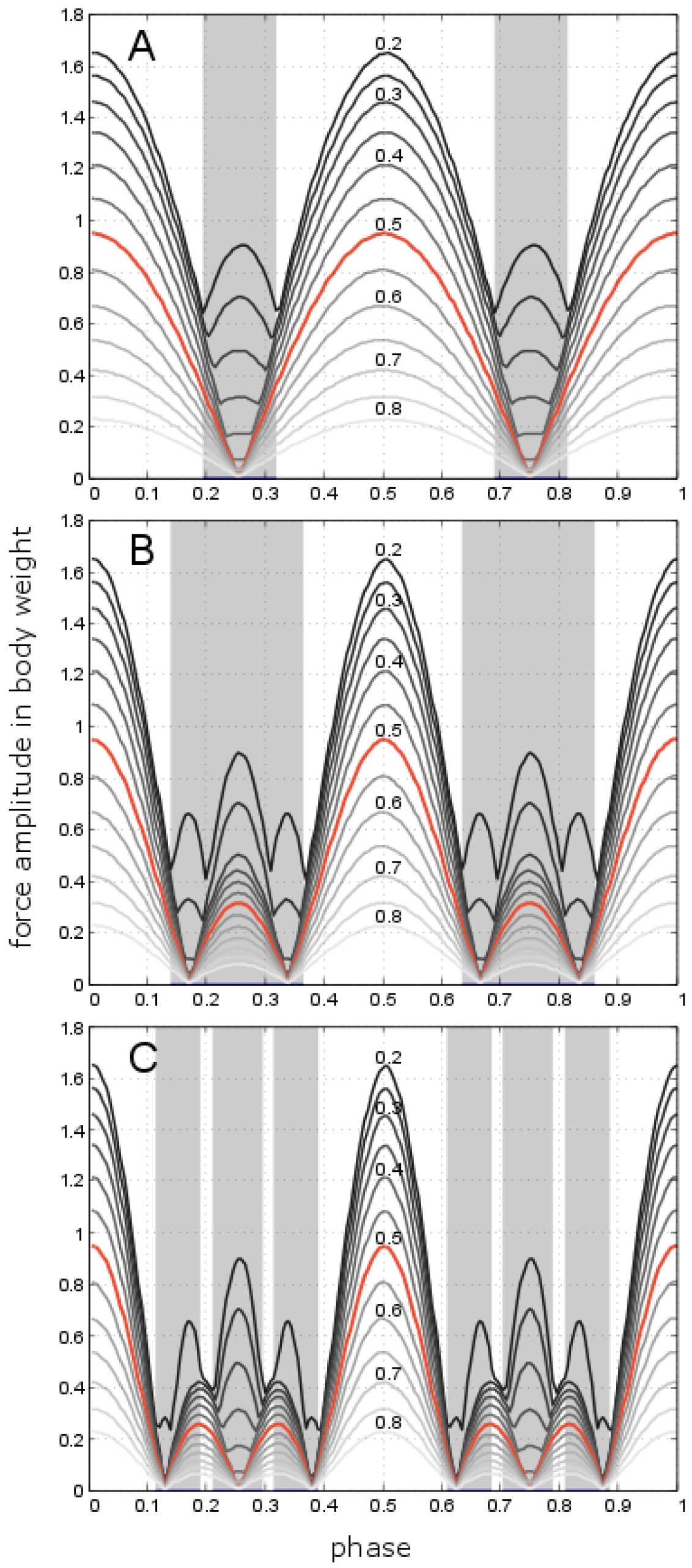
Dependency of total leg force amplitudes on the phase shift of ipsilateral adjacent legs and duty factor for locomotor apparatuses with two to four leg pairs. A) Two leg pairs: Duty factors range from 0.2 (dark grey) to 0.8 (light grey) with the duty factor of 0.5 being highlighted in red. With low duty factors and intermediate phase shifts the peak frequency of the force oscillations deviates from two times stride frequency; these Intervals are shaded in grey. B) Three leg pairs; C) Four leg pairs.

In locomotor apparatuses with three pairs of legs, vertical ground forces maintain only half of their maximum amplitude if θ is 0.4 (Fig. 3). The phase shift at which amplitude values are halved is independent of the duty factor, but changes with the number of legs and adopts a saturation curve (Fig. 4). Thus, the halving phase shift is 0.427 in locomotor apparatuses with four pairs of legs and 0.443 in those with 5 pairs of legs. In locomotor systems with higher leg numbers, values progressively converge to 0.5. However, with only two pairs of active walking legs the critical phase shift is 0.34, representing the highest difference from the alternating case. The force amplitude minima show a similar behaviour to the half peak values with respect to θ (Fig. 4). In locomotor apparatuses with three pairs of legs, minimum force amplitudes occur at θ = 0.34; with four pairs of legs it is 0.38; and with 5 pairs of legs it is 0.4. With more pairs of legs, minimum force amplitudes occur at even larger phase shifts, i.e. smaller deviations from θ = 0.5. Locomotor apparatuses with only two pairs of legs undergo relatively high force amplitudes in the phase range from 0.2 and 0.3 if duty factors are low (Fig. 2, Fig. 4); therefore, the position of the force amplitude minimum changes with the duty factor. The values are as high as 0.32 with a duty factor of 0.3 and decrease towards 0.25 at higher duty factors.

**Fig. 3.**
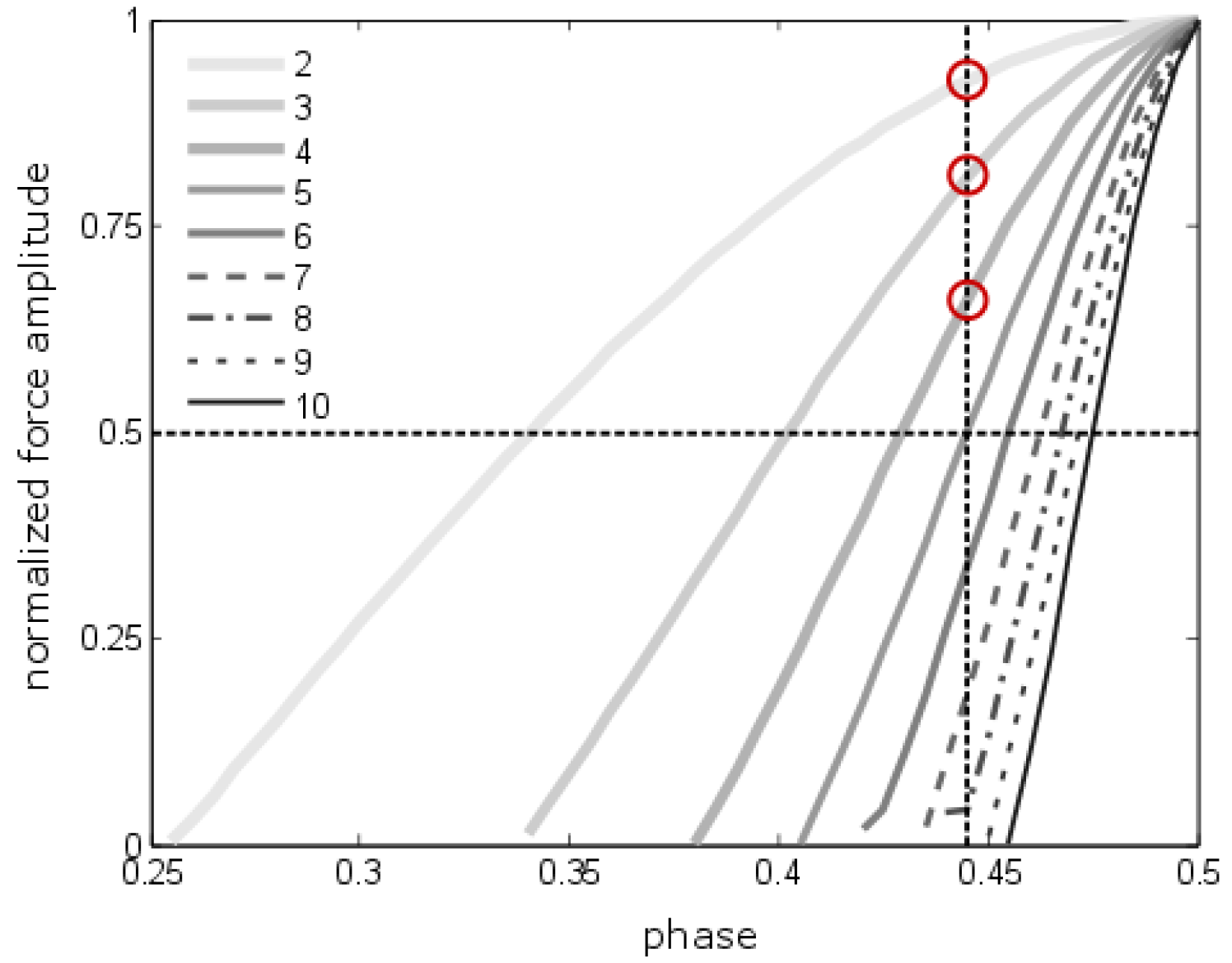
Normalised amplitudes of the overall vertical ground forces for locomotor apparatuses with two to ten pairs of legs (legend) and a duty factor of 0.5 between the maximum at alternating sets of legs and the first minimum (cp. Fig. 2). The slope of the amplitudes increases significantly with an increasing number of leg pairs. The trajectories’ points of intersections with the horizontal dashed line indicate those θ at which only ½ of the maximum force amplitude is retained (see Fig. 4). The vertical dashed line represents a deviation of 0.06 from the strictly alternating gait pattern as found in fast running desert ants (see main text for further explanations). The red circles indicate the force amplitude values for locomotor systems with two, three and four leg pairs at such a deviation from strict alternation.

**Fig. 4.**
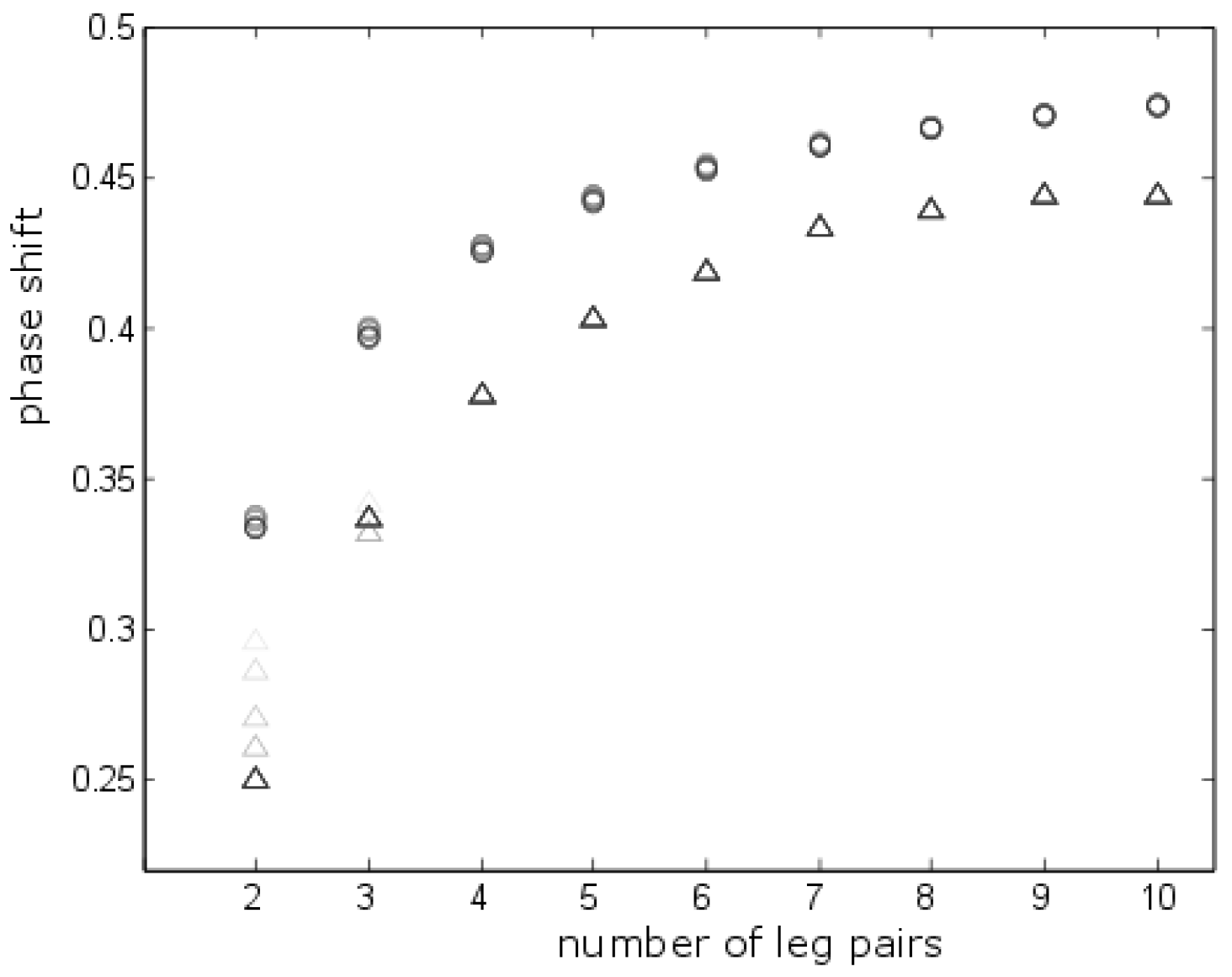
Position of the first amplitude minimum (triangles) and half peak values (circles) with respect to θ for locomotor apparatuses with two to ten pairs of walking legs. Particularly for low leg numbers the position of the force amplitude minimum decreases with increasing duty factors (decreasing brightness, cp. Fig. 2A) while duty factors have virtually no impact on the phase shift at which force amplitudes are halved (see Figs. 2, 4).

Specifically at low duty factors and intermediate ipsilateral phases, harmonics may occur that result in oscillation frequencies of the vertical forces adopting multiples of two-times stride frequency (i.e. the value to be expected for alternating leg coordination and required for appropriate loading of leg springs). Phases with oscillation frequencies higher than two range: between 0.2 and 0.32 in locomotor apparatuses with two pairs of legs; between 0.14 and 0.36 in locomotor apparatuses with three pairs of legs; and between 0.12 and 0.39 in those with four pairs of legs (Fig. 2).

Ipsilateral phase shifts significantly deviating from 0.5 result in an increase of the minimum number of legs in contact with the ground. Sufficiently high deviations can even prevent phases without any legs on the ground, although single leg duty factors are then significantly lower than 0.5 (Fig. 5).

**Fig. 5.**
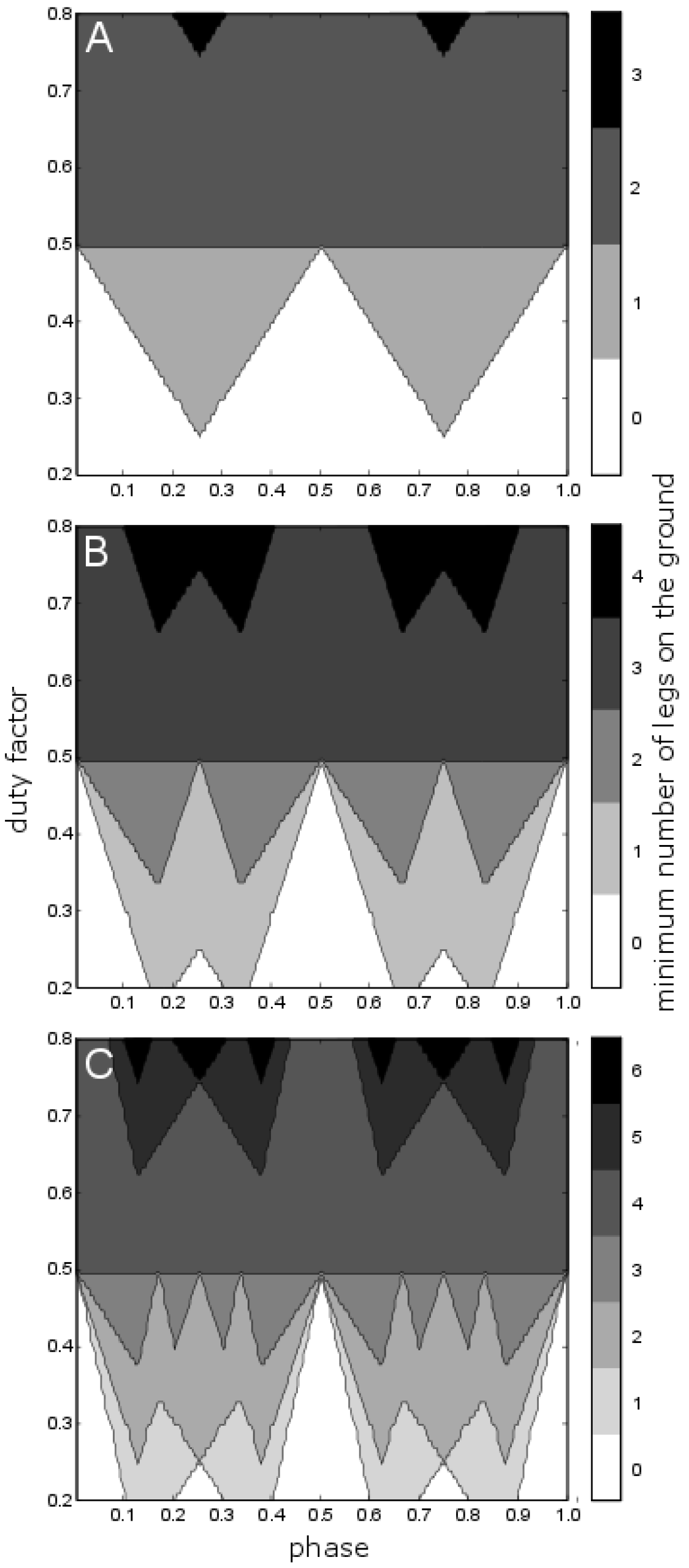
Minimum number of legs on the ground as function of duty factor and phase shift of ipsilateral adjacent legs. A) Two pairs of legs B) Three pairs of legs C) Four pairs of legs. With sets of legs close to alternating activity and duty factors below 0.5 the minimum number of legs on the ground is always 396 0 while phase shifts deviating from alternation (0.5) and synchrony (0 and 1) result in increased minimum numbers of legs in contact with the ground.

## Discussion

The efficiency of bouncing gaits such as running, trotting or the alternating tripodal gait of insects and harvestmen largely depends on elastic structures in the walking legs. Since stress-strain dependencies of tendons and equivalent structures are mostly nonlinear and characterised by significant hysteresis (Bennet et al., 1986; Blickhan, 1986; Dudek and Full, 2006), these springs have to undergo sufficient strain in order to be useful as energy storage, which requires sufficient load during the legs’ stance phase (Rubenson et al., 2004). In steady locomotion leg loading is dominated by the vertical force components. The changes of these forces result in length changes of elastic structures within the legs and can therefore be used for cyclic energy storage and recovery.

However, just as temporally largely overlapping ground contacts decrease force amplitudes and may prevent sufficient loading of elastic structures, the use of spring mass dynamics may also be hampered if loading forces exceed the physical abilities of the legs’ musculoskeletal structure (Seyfarth et al., 2001; Taylor and Dirks, 2012). Accordingly, in horses the change from fast trot to slow gallop minimizes the peak values of total vertical forces (Farley and Taylor, 1991) and single leg forces drop slightly (Biewener and Taylor, 1986). Interestingly in many species the change from trot to gallop seems to coincide also with an optimization of metabolic costs (Hoyt and Taylor, 1981; Minetti et al., 1999). In sideways running ghost crabs, however, such an energy optimisation could not be validated (Full and Weinstein, 1992), which might relate to a lack of elastic energy recovery during gallop.

In gaits with strictly alternating leg coordination, maximum vertical ground forces and duty factors depend on each other. While duty factors and contact duration decline with increasing running speed, peak forces increase in order to keep the momentum constant (Blickhan, 1989). If leg synchronisation declines, running speed can be increased and the duty factors can decline without significant increases in total vertical peak forces (Fig. 1). With declining duty factors, and a relative increase of coordination errors as found in fast running desert ants (Wahl et al., 2015) or on rough terrain (Sponberg and Full, 2008), with alternating leg coordination and high vertical amplitudes of the COM the animals are at risk of touching down initially only with one leg. Resulting high single leg forces might exceed the physical properties of the leg adjusted to shared force application as typical for straight, level locomotion. Overload, in turn, can cause a failure of the locomotor apparatus. Metachronal leg coordination - i.e. coordination patterns with ipsilateral phase shifts significantly deviating from those found in bouncing gaits - results in minimised total force maxima (Fig. 2). Accordingly, desynchronization of the sets of legs also minimises the maximum feasible load on a single leg and thereby may help to avoid such collapses. Accordingly, blaberid cockroaches change from an alternating tripodal leg coordination scheme at intermediate running speeds to metachronal leg coordination at maximum speeds (Weihmann et al., 2017), preventing large vertical amplitudes of the COM and vertical force peaks. Nevertheless, some arthropod species specialized for high speed escape runs are able to move even bi-pedally using specifically adjusted legs when extremely high speeds are required (Burrows and Hoyle, 1973; Full and Tu, 1991).

With increasing running speed, given leg lengths require decreasing contact periods and quicker actuation of the legs during stance. At a higher contraction speed, muscular forces decrease and the required leg forces can be maintained only by increased consumption of metabolic energy (Hill, 1938; Hill, 1939). In bouncing gaits, increasing running speeds result in increasing relative swing phase durations and decreasing duty factors. The synchronised leg activity leads to phases without any contact with the ground if duty factors fall below 0.5. Metachronal coordination of ipsilateral legs enables increasing relative swing phase durations without ballistic phases and with reduced vertical peak forces. Accordingly, in both cases, muscle activity during swing can be slower and more energy efficient (Garcia et al., 2000; Marsh et al., 2004; Nishii, 2006) which may trigger reduced metabolic costs as found in fast moving desert beetles (Bartholomew et al., 1985).

Even in specialist high speed runners the maintenance of synchronicity among the legs of the alternating sets seems to be challenged at high running speeds. Though contact duration typically decreases exponentially with speed (Blickhan et al., 2005; Reinhardt and Blickhan, 2014; Weihmann, 2013), the tripod synchrony factor, i.e. the normalized fraction of contact phase overlap between legs in the same set of legs (Spagna et al., 2011), does not exceed 0.8 in fast running desert ants of the species *Cataglyphis fortis* (Wahl et al., 2015). Accordingly, with increasing running speeds the impact of internal and external disturbances seems to increase too. Decreased synchronisation however results in reduced amplitudes of total vertical forces, lower vertical amplitudes of the COM and a lower capacity for elastic energy storage in the legs.

The number of active pairs of legs significantly impacts the requirements for coordination accuracy. While in locomotor systems with three pairs of legs a phase deviation of 0.1 from the alternating pattern results in halving of the total vertical peak forces, this occurs at deviations of only 0.07 in systems with four pairs of legs and at even smaller deviations in locomotor systems with more pairs of legs (Fig. 4). Assuming a duty factor of 0.3, a tripod synchrony factor of 0.8 - as found in fast running *C. fortis* (Wahl et al., 2015) - already leads to phase shifts of approximately 0.3-(0.3-0.8)=0.06 away from the strict alternating pattern. Such a shift reduces the ants’ vertical force amplitudes by 19% (Fig. 3). In locomotor systems with four pairs of legs however, the same shift would result in a reduction by a third. Quadrupeds in turn can cope with higher phase variations since phase deviations of 0.06 reduces peak forces by only 7%. Accordingly, in animals with fewer legs, ipsilateral phase shifts can be relatively large without causing significant reductions in total vertical peak forces, vertical amplitudes of the COM and the capacity for elastic energy storage. Animals with more than about five pairs of active legs, even fast species such as centipedes (Lewis, 1981), are practically incapable of employing spring mass dynamics in the sagittal plane. In such species, the tiny inaccuracies induced by internal and external perturbations prevent sufficient synchronisation of the putative sets of diagonally adjacent legs.

Many advantages of spring mass dynamics as found in running and trotting gaits, such as energy efficiency and running stability (Blickhan et al., 2007), are fully effective only on relatively firm and smooth substrates and require specialized locomotor apparatuses enabling transient elastic energy storage. For many species that lack such specific adaptations, such as spiders (Sensenig and Shultz, 2003; Weihmann, 2013), many bugs, beetles (Goyens et al., 2015), amphibians (Ahn et al., 2004) or tortoises (Zani et al., 2005), disadvantages of bouncing COM trajectories may prevail. Increased peak forces due to constructive interference of single leg ground forces, may then rather be a perturbation of movement stability and vision (Ardin et al., 2015; Zurek and Gilbert, 2014) which are avoided by high duty factors and intermediate θ.

High synchronised leg forces are also transferred onto the substrate which may transmit the elicited vibrations to potential predators or prey organisms (Barth, 2002; Magal et al., 2000). The temporally distributed force application of gait patterns with intermediate θ in turn may camouflage the signals and prevent the assessment of the producer’s location, speed and size. Granular substrates can in turn dissipate significant proportions of movement energy - in particular if peak forces are high - largely preventing energy recovery (Gravish et al., 2014; Hafemann and Hubbard, 1969; Le Jeune et al., 1998). This external energy dissipation also makes lower speeds with resulting high duty factors and intermediate θ preferable.

Close to and between the positions of the force amplitude minima (i.e. between 0.14 and 0.36 in hexa-pedal locomotor systems), the peak frequency of the force oscillations differs significantly from two times stride frequency; specifically at low duty factors (Fig. 2). Harmonics occurring in this range lead to multiples of the value expected for spring mass dynamics. Accordingly, within this range COM oscillations cannot be effectively used for the initiation and maintenance of symmetrical bouncing gaits. In horses typical gaits using ipsilateral phase shifts in the range of minimum vertical COM amplitudes and high oscillation frequencies are ambling gaits such as tölt, single-foot and rack (Hildebrand, 1989; Robilliard et al., 2007). The equivalent in hexapedes is the high speed metachronal gait as found in cockroaches and mites (Weihmann et al., 2017; Weihmann et al., 2015). Since contralateral phase shifts are not considered the present model is not suitable for the examination of asymmetric gaits such as gallop (Hildebrand, 1965). For gait patterns as found in mammals galloping at high speeds (Biancardi and Minetti, 2012) in addition to ipsilateral phase shifts of about 0.3 contralateral phase shifts lower than 0.3 are required. Such leg coordination patterns, however, would re-establish bouncing dynamics of the COM at which the sequenced and overlapping ground contacts of the four legs substitute a single leg ground contact of the traditional spring-mass model (Bertram and Gutmann, 2009; Ruina et al., 2005). The prime example for galloping arthropods are semi-terrestrial sideways moving ghost crabs that reach impressive speeds in their typical flat sand habitats in the intertidal zone (Full and Weinstein, 1992). Among fully terrestrial fast moving arthropods however, so far, no such contralateral phase adaptations have been found except for a brief notice referring to the peculiar high speed jumping gait of the jumping bristletail *Petrobius* (Manton, 1972).

Ipsilateral phase shifts that differ significantly from 0.5 result in an increased minimum number of legs on the ground (Fig. 5). Sufficiently high deviations from θ = 0.5 completely prevent airborne intervals even though duty factors are considerably lower than 0.5. Consequently, such gaits facilitate permanent proprioceptive feedback and may increase controllability and stability of the locomotion.

Apparently, locomotor systems of bipeds with only one pair of walking legs largely avoid force interference problems in particular at high running speeds (Alexander and Jayes, 1978; Geyer et al., 2006). Consequently also perturbations caused by interfering legs may not occur and the stability of COM dynamics and the coordination patterns should be higher. In this regard, the control effort (Haeufle et al., 2014) is expected to be much lower than in poly-pedal designs, which probably outweigh reduced static stability and may represent a major advantage of bipedal body plans.

## Methods and Procedures

Changes of ipsilateral phase shifts and changed duty factors were examined with regard to total vertical forces for locomotor systems with two to ten pairs of walking legs. All analyses were performed using MatLab scripts (MATLAB 7.10.0; The MathWorks, Natick, MA, USA). Since the study is focused on general understanding of poly-pedal locomotion specific anatomies of walking legs have not been taken into regard.

Phase relations between the legs were defined with regard to stride durations which were set to 1. The phase shift θ was determined by the occurrence of one leg’s touchdown within the stride period of another leg; it was always between 0 and 1. Thus, the phase shift was 0.2 if the touchdown of the second leg occurred after 20% of the stride period of the first one. Since phase relations follow a circular distribution phase shifts of 0 and 1 are equivalent (Weihmann, 2013). The analyses went through all θ from 0 to 1; duty factors from 0.2 to 0.8 were examined, i.e. the study focused on physiologically relevant values.

In previous studies it has been shown that the contralateral phase relation between the rear legs of cockroaches and mites is 0.5 irrespective of running speed (Weihmann et al., 2017; Weihmann et al., 2015). Accordingly and in order to reduce the number of variables, contralateral phase relations between the rear legs were set to 0.5, i.e. rear legs alternated strictly. Setting contralateral phases of the rear legs to 0.5 restricts the approach to symmetrical (Hildebrand, 1965) gait patterns. During straight level terrestrial locomotion the vertical ground force component is always the largest, irrespective of the number of legs used or which class of animals is taken into account (Biewener, 2003). Moreover, vertical force components are most decisive and characteristic for COM dynamics of certain gaits even though lateral forces can reach significant values in animals with sprawled legs (Blickhan and Full, 1993; Reinhardt et al., 2009). Accordingly the present study is focussed on vertical forces. As a first approximation the force applied during the stance phase of a leg has been modelled as 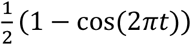 for the time interval *t* from 0 to 1, i.e. the start and the end of the stance phase (cp. Fig. 1). Each stride is comprised of an initial stance and a subsequent swing phase where no forces are applied onto the ground. The quotient of contact duration and stride duration defines the duty factor of a leg. Stride duration, duty factors and ground forces were equal for all legs. In order to generate different gait patterns ipsilateral leg activities were shifted against each other according to prescribed θ from 0 to 1. Ipsilateral phase, duty factor and leg number were the only changed parameters; oscillations of total vertical forces, then, emerged from the interplay of these kinematic measures and the stereotyped single leg ground forces.

With strictly alternating sets of legs overall ground forces over a stride have two maxima which are caused by the consecutive activity of the sets of legs (Fig. 1E). All forces were summed up and divided by the mean value resulting in normalization onto body weight. Spectral analyses of the resulting force oscillations were accomplished by using the Fast Fourier Transformation algorithm implemented in Matlab. Resulting amplitudes correspond to 1/2 of the peak to peak values of the respective ground forces, i.e. the difference between the absolute maximum and minimum values of summed overall ground forces. Amplitude values change symmetrical about the alternating pattern (θ = 0.5). Accordingly amplitude values for θ = 0.4 and θ = 0.6 are equivalent. In the results section, therefore, values for local minima, half peak values etc. are provided only for the phase range between 0 and 0.5.

The minimum number of legs in contact with the ground can serve as proxy for the requirement of dynamic stabilization of a gait (Ting et al., 1994). The number of legs on the ground was determined equivalent to the ground forces by summing up the legs in contact with the ground for each instant of time; minimum values were determined for the increments of θ and duty factor subsequently.

## Data availability

All relevant data and maths are provided within the manuscript and figures.

## Authors’ contributions

n/a

## Research Ethics

n/a

## Animal ethics

n/a

## Permission to carry out fieldwork

n/a

## Acknowledgement

I would like to thank Michael Günther and Walter Federle for inspiring discussions. I thank Silvia Daun and Ansgar Büschges for their critical reviews and Sam Skeaping and Emily Pycroft for copyediting the manuscript.

